# The renin angiotensin system in cognitive flexibility: Effects of single-dose losartan on task-switching in healthy volunteers

**DOI:** 10.1101/2025.11.23.689943

**Authors:** Divya Prasad, Georgia Feltham, Riccardo De Giorgi, Sara Costi, Elaine Fox, Amy L. Gillespie, Andrea Reinecke

## Abstract

**Introduction:** Cognitive flexibility, defined as the ability to shift mental strategies when environmental demands change, may influence psychological well-being and psychiatric treatment outcomes. Emerging research increasingly implicates the renin angiotensin system (RAS) in cognition, yet its role in cognitive flexibility is inadequately understood.

**Methods:** We conducted a double-masked trial with N=60 healthy adults aged 18-50. Participants were randomly assigned to receive a single 50 mg dose of losartan, an angiotensin II receptor blocker, or placebo. Following a waiting period, participants completed a task-switching paradigm as a measure of cognitive flexibility, which required dynamically categorizing arrows based on location or direction. Statistical analyses were conducted to determine group (losartan vs. placebo), task (location vs. direction), and switch (switching vs. repeating a task) effects on reaction time (RT) and accuracy independently. A composite bin score was also computed, integrating both RT and accuracy data.

**Results:** All participants displayed the expected switch costs. While participants on losartan exhibited numerically superior RTs and accuracy, losartan did not significantly improve task-switching performance relative to placebo. This was confirmed by the bin score analysis, which indicated no benefit of losartan when RT and accuracy were considered together. An exploratory probe suggested that losartan improved accuracy on direction switch trials, indicating a potential drug effect under specific cognitive conditions.

**Conclusion:** Overall, we found no significant benefits of losartan on task-switching in healthy adults. Our results may be explained by ceiling effects in our sample, which could have hindered the detection of group differences, or possibly reflect a negligible role for the RAS in cognitive flexibility.

## Introduction

To survive and thrive, humans must switch between different mental strategies as environmental demands evolve. This cognitive flexibility (Dajani & Uddin, 2015) allows us to attain diverse goals in a dynamic world and is associated with several positive outcomes, including greater stress resilience (Genet & Siemer, 2011), quality of life (Davis et al. 2010), and post-traumatic growth (Hijazi, Keith, & O’Brien, 2015). Conversely, cognitive inflexibility is linked to various psychiatric symptoms, including anxiety (Wilson et al. 2018; Rosa-Alcázar et al. 2020; Gustavson et al. 2017; Johnco, Wuthrich, & Rapee, 2015), depression (Murphy et al. 2012; Maramis et al. 2021; Yu, Yu, & Lin, 2020; Johnco, Wuthrich, Rapee, 2015), post-traumatic stress disorder (PTSD; Keith, Velezmoro, & O’Brien, 2015; George et al. 2015), obsessive compulsive disorder (OCD; Rosa-Alcázar et al. 2020; Gruner & Pittenger, 2017; Rosa-Alcázar et al. 2020), and borderline personality disorder (Aslan, Grant, & Chamberlain, 2023; Duda et al. 2024). Cognitive flexibility may also influence treatment outcomes (Johnco, Wuthrich, & Rapee, 2014; Nagata et al. 2018; Deveci et al. 2023; Jalal et al. 2018; Ben-Zion et al. 2018) and facilitate better application of psychotherapeutic strategies (Johnco, Wuthrich, & Rapee, 2014). Further, research suggests that survivors with higher cognitive flexibility 1-month post trauma show less severe PTSD symptoms 13 months later (Ben-Zion et al. 2018). Together, this evidence implicates cognitive flexibility in psychological well-being and interventional response, whereas cognitive inflexibility may increase vulnerability to and/or maintain psychopathology. Thus, there is scientific and clinical rationale to elucidate the underpinnings of cognitive flexibility and develop novel strategies to enhance it.

The renin angiotensin system (RAS), traditionally known for its role in blood pressure regulation, has been increasingly implicated in cognition (Zhou et al. 2024; Chrissobolis et al. 2020). For instance, a single dose of the angiotensin II receptor blocker (ARB) losartan improves prospective memory (Mechaeil et al. 2011), promotes the extinction of fear memories (Zhou et al. 2019), biases learning toward positive instead of negative events (Pulcu et al. 2019) and enhances mnemonic discrimination in healthy volunteers (Prasad et al. 2025). While these processes may rely on or facilitate flexible thinking, little research has examined RAS influence on cognitive flexibility directly.

Studies have reported mixed results surrounding ARB effects on cognitive flexibility (Hajjar et al. 2022; Ho & Nation, 2017; Saxby et al. 2008; Wharton et al. 2015; Hajjar et al. 2020). For instance, some evidence suggests that ARB treatment may improve or preserve set-shifting abilities in older adults with and without mild cognitive impairment (Hajjar et al. 2022; Ho & Nation, 2017). Yet, other research suggests no significant effect of ARB treatment on set-shifting in older adults with hypertension, despite other cognitive benefits (Saxby et al. 2008). The relationship between the RAS and cognitive flexibility remains unclear, prompting further exploration for several reasons. First, it is unknown whether the RAS affects cognitive flexibility in adults without cognitive impairment and in earlier stages of life, when the onset and functional burden of mental illness may be particularly salient (Kessler et al. 2005; Solmi et al. 2022). This would represent a fundamental step in determining any baseline augmentative effects of ARBs on cognitive flexibility, which may then be harnessed to restore deficits preventatively or curatively in psychiatric populations. Second, it would be useful to probe RAS influence on more effortful forms of cognitive flexibility, which is currently lacking given the simplicity of previous paradigms (Bunge & Zelazo, 2006; Saxby et al. 2008; Hajjar et al. 2022; Ho & Nation, 2017). To address these gaps, we conducted a double-masked experimental medicine trial examining the effects of acute losartan on task-switching in healthy adults. Given previous observations of ARB-mediated benefits to cognitive mechanisms (Mechaeil et al. 2011; Zhou et al. 2019; Pulcu et al. 2019; Prasad et al. 2025) and set-shifting (Hajjar et al. 2022; Ho & Nation, 2017), we hypothesized that losartan would reduce RT and accuracy switch costs, reflecting enhanced cognitive flexibility.

## 2 Methods

### 2.1 Sample

This trial was approved by the University of Oxford research ethics committee (CUREC: R80447). N=60 healthy participants aged 18-50 years were recruited through university notice boards and social media. Sample size was informed by the only cognitive losartan study in healthy humans available at the time (ηp^2^ = 0.13; Mechaeil et al., 2011; power analysis conducted in G*Power for a repeated-measures ANOVA, within-subject interaction, F = 0.15, β = 0.80, α = 0.05; Faul, Erdfelder, Lang, & Buchner, 2007).

All participants provided written informed consent prior to the testing visit and did not meet criteria for any DSM-5 diagnosis, had not taken psychoactive drugs in the 6 weeks prior to testing, had a BMI of 18-30, and were non- or light-smokers (<5 cigarettes a day). All participants completed “Spot-the-Word” (STW) as a measure of verbal intelligence (Baddeley et al. 1993).

### 2.2 Procedures

Participants completed self-report questionnaires evaluating trait anxiety, depression, and other psychological parameters (see Supplementary Material). Participants were then randomly administered 50 mg oral losartan (Cozaar; Merck Sharp & Dohme Ltd., Hertfordshire, UK) or matched placebo, according to a 1:1 allocation ratio. Randomization was stratified by sex assigned at birth.

Following a 1-hour waiting period, when losartan levels peak in plasma (Lo et al. 1995), participants completed a neurocognitive task battery, featuring an asymmetric task-switching paradigm (Gustavson et al. 2017). Task-switching paradigms have been widely used to probe cognitive flexibility, as they require intermittent modification of a response strategy based on changing cues (Todorovic et al. 2022; Dajani & Uddin, 2015; Monsell, 2003). Subjects are required to repeat a previously applied strategy or switch to a different one, typically displaying faster RTs for repeat vs. switch conditions and consequently, a ‘switch cost’. In the present study, participants were instructed to categorize arrow stimuli (<<<<< or >>>>>) based on a changing cue (+ or *). When a + was shown, participants were required to respond with where the arrows were on the screen (location task), and when a * was shown, they were required to respond with the direction the arrows were facing (direction task). On each trial, the cue appeared for 750ms, indicating which task to complete. To mitigate participants responding correctly at random or if they misunderstood, locations and directions of the arrows were incongruent on most trials (e.g., if the stimulus was shown on the right side of the screen, it pointed left 75% of the time). Participants began by completing practice blocks for each task and were then tested on four experimental blocks with mixed task demands. These blocks presented the cues in random order, with 112 trials per block and 448 trials in total. Within each block, participants completed 28 switch trials, 16 1-repeat trials, 8 2-repeat trials, and 4 3-repeat trials (56 for location and 56 for direction; Gustavson et al. 2017).

Heart rate and blood pressure were also assessed prior to drug administration and at peak-level. After testing, participants and experimenters guessed which treatment had been administered to determine whether masking was successful.

### 2.3 Outcomes and statistical analysis

The analysis plan was pre-registered on AsPredicted.org (#168742). Data cleaning and statistical analyses were conducted using R Version 4.1.1. Changes in heart rate, systolic, and diastolic blood pressure from baseline to drug peak level were investigated using repeated measures ANOVAs with time as the within-subjects factor (pre- or post-administration) and group as the between-subjects factor (losartan or placebo).

Task data were processed based on previous studies (Wilcox & Keselman, 2003; Gustavson et al. 2017; Todorovic et al. 2022; see Supplement A). Two repeated-measures ANOVAs were conducted, where group (losartan, placebo), task (location, direction), and switch condition (repeat, switch) were factors. RT (ms) and accuracy (%) were dependent variables. We further explored losartan’s effect on switch costs through a composite bin scoring method which integrated RT and accuracy (Hughes et al. 2014; Draheim et al. 2016). For each participant, average repeat trial RTs were calculated and subtracted from each of their switch trial RTs. Then, the RT differences on correct trials were rank ordered into 10 deciles and accordingly scored from 1-10. Incorrect trials were assigned a bin value of 20. All bin values were then summed per participant and averaged across participants in each treatment group. Group bin scores were then compared via an independent samples t-test, with a lower bin score indicating better task-switching performance.

## 3 Results

### 3.1 Participant Characteristics and Drug Effects

After outlier exclusion due to low accuracy, the final sample consisted of N=58 participants (N=30 losartan, N=28 placebo), who were well-balanced on sociodemographic and clinical parameters and verbal intelligence (Table 1). No acute drug effects on heart rate, systolic, or diastolic blood pressure (all ps>0.21) were observed. Neither experimenter nor participants guessed drug administration above chance level (correct participant = 39%, correct experimenter = 49%, both ps>.11), indicating that masking was maintained. There was minimal data loss (1 STW score, 1 experimenter/participant allocation guess).

**Table 1:** A./B. Sociodemographic, clinical, and personality characteristics of participants in the two groups (means and standard deviations). C. Cardiovascular markers at baseline and at drug-peak level (means and standard deviations).

| Measure | Losartan (n = 30) | Placebo (n = 28) |
| --- | --- | --- |
| **A. Sociodemographic Data** |  |  |
| Sex assigned at birth (Female/Male), n | 20 / 10 | 19 / 9 |
| Age, years | 25.70 (6.18) | 27.14 (8.28) |
| Education, years | 17.67 (2.66) | 17.91 (2.28) |
| Verbal IQ (Spot-the-Word) | 70.76 (11.63) | 72.25 (11.28) |
| **B. Clinical and Personality Measures** |  |  |
| State Trait Anxiety, STAI | 37.37 (8.20) | 35.29 (8.82) |
| Depressive Symptoms, BDI | 3.93 (4.40) | 5.79 (6.85) |
| Attentional Control Scale, ACS | 58.53 (6.07) | 58.46 (6.85) |
| Intolerance of Uncertainty Scale, IUS | 54.27 (14.13) | 50.82 (13.46) |
| Perceived Stress Scale, PSS | 20.50 (6.28) | 20.61 (6.93) |
| Temporal Experience of Pleasure Scale, TEPS | 85.10 (8.89) | 86.90 (8.60) |
| **C. Heart Rate and Blood Pressure (BP)** |  |  |
| Heart Rate before administration | 73.33 (11.73) | 76.82 (15.25) |
| Heart Rate after administration | 63.87 (9.76) | 66.14 (10.92) |
| Systolic BP before administration | 112.67 (9.97) | 111.82 (13.28) |
| Systolic BP after administration | 112.23 (9.39) | 108.50 (9.58) |
| Diastolic BP before administration | 71.03 (5.75) | 74.29 (9.62) |
| Diastolic BP after administration | 70.90 (7.53) | 71.79 (8.70) |

### 3.2 Reaction Time Analyses

We found significant main effects of task (F(1, 56)=131.32, p<0.001, ηp^2^ = 0.70) and switch condition (F(1, 56)=17.11, p<0.001, ηp^2^=0.23) on reaction time (see Figure 1). Across groups, participants were slower to indicate where the arrows were pointing (direction task) compared to where the arrows were located on the screen (location task), as expected. Participants were also slower to switch tasks than repeat them, replicating previously observed switch costs (Monsell, 2003; Gustavson et al. 2017; Todorovic et al. 2022). A significant task*switch interaction was observed (F(1, 56)=6.50, p=0.014, ηp^2^= 0.10): participants demonstrated greater switch costs when switching from the harder to the easier task (location switch, 32.8ms) than when switching from the easier to the harder task (direction switch, 7.3ms), aligning with previously reported asymmetric switch costs (Todorovic et al. 2022; Gustavson et al. 2017). No significant main or interaction effects of group emerged (all ps>0.243), indicating that losartan and placebo groups responded with comparable speed (Figure 1).

**Figure 1:**
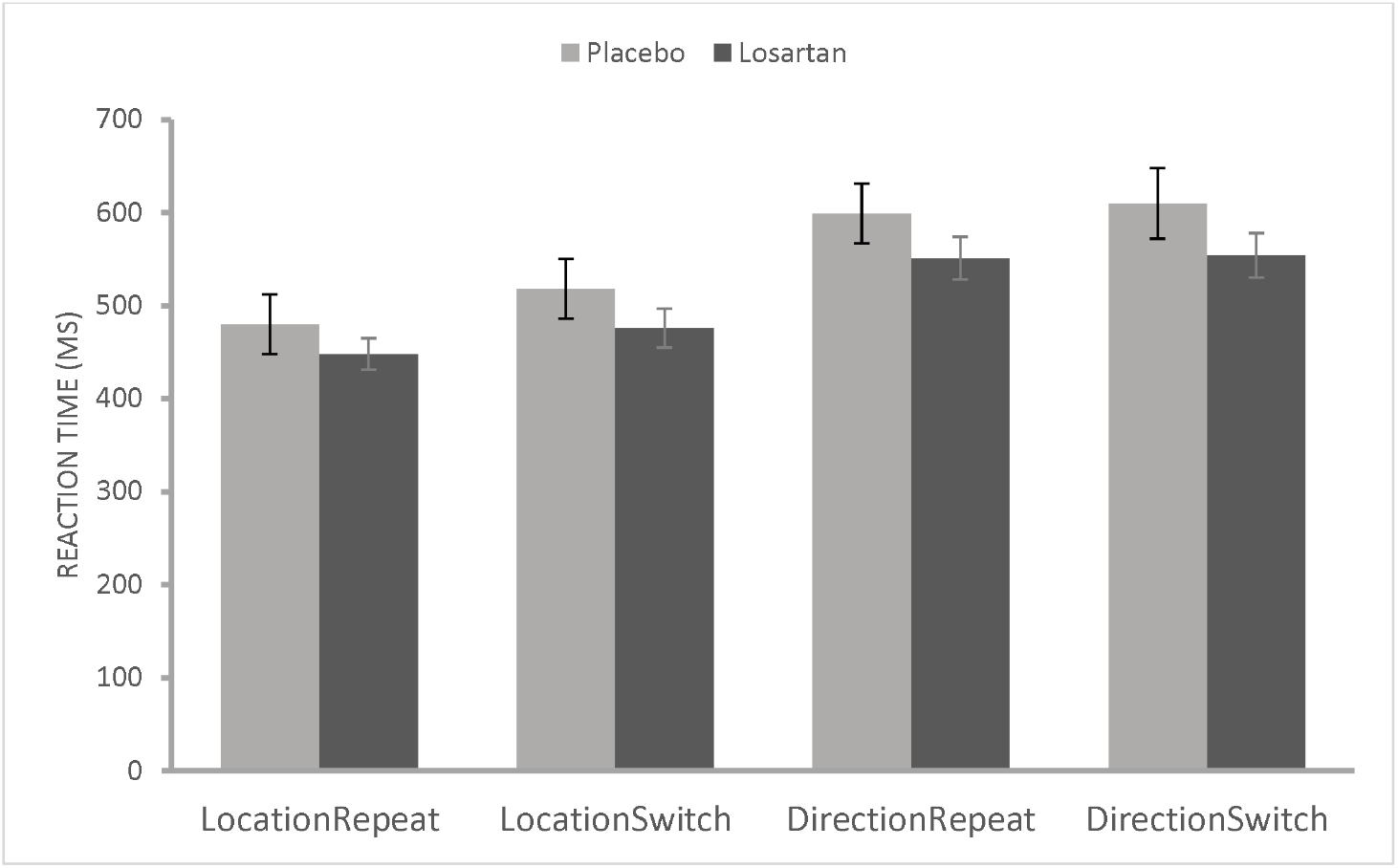
Mean RTs by task (location/direction) and switch condition (repeat/switch) across losartan and placebo groups, on correct trials only.

### 3.3 Accuracy

Similarly, we found significant main effects of task (F(1, 56)=4.68, p=0.04, ηp^2^= 0.07) and switch condition (F(1, 56)=10.81, p=0.002, ηp^2^=0.16) on accuracy (see Figure 2). Contrary to our expectation, participants were more accurate to say which way the arrows were pointing (direction task) compared to where the arrows were on the screen (location task). As expected however, participants were less accurate on switch trials versus repeat trials, demonstrating an accuracy switch cost. We also observed a significant task*switch interaction (F(1, 56)=16.86, p<0.001, ηp^2^=0.23) such that participants showed greater accuracy costs when switching from direction to location than the other way, demonstrating switch cost asymmetry (Gustavson et al. 2017). No significant main or interaction effect of group emerged, though the group main effect trended toward significance (F(1, 56) = 3.23, p=0.078, ηp^2^=0.05). Exploratory pairwise comparisons of estimated marginal means showed a significant group effect on direction-switch trials, p =.031 (Bonferroni-adjusted), Cohen’s d=0.59, such that the losartan group was more accurate than the placebo group when switching from location to direction (mean difference= 0.022, SE = 0.010, 95 % CI [0.002, 0.041]). No other conditions showed significant group effects (all ps > .07).

**Figure 2:**
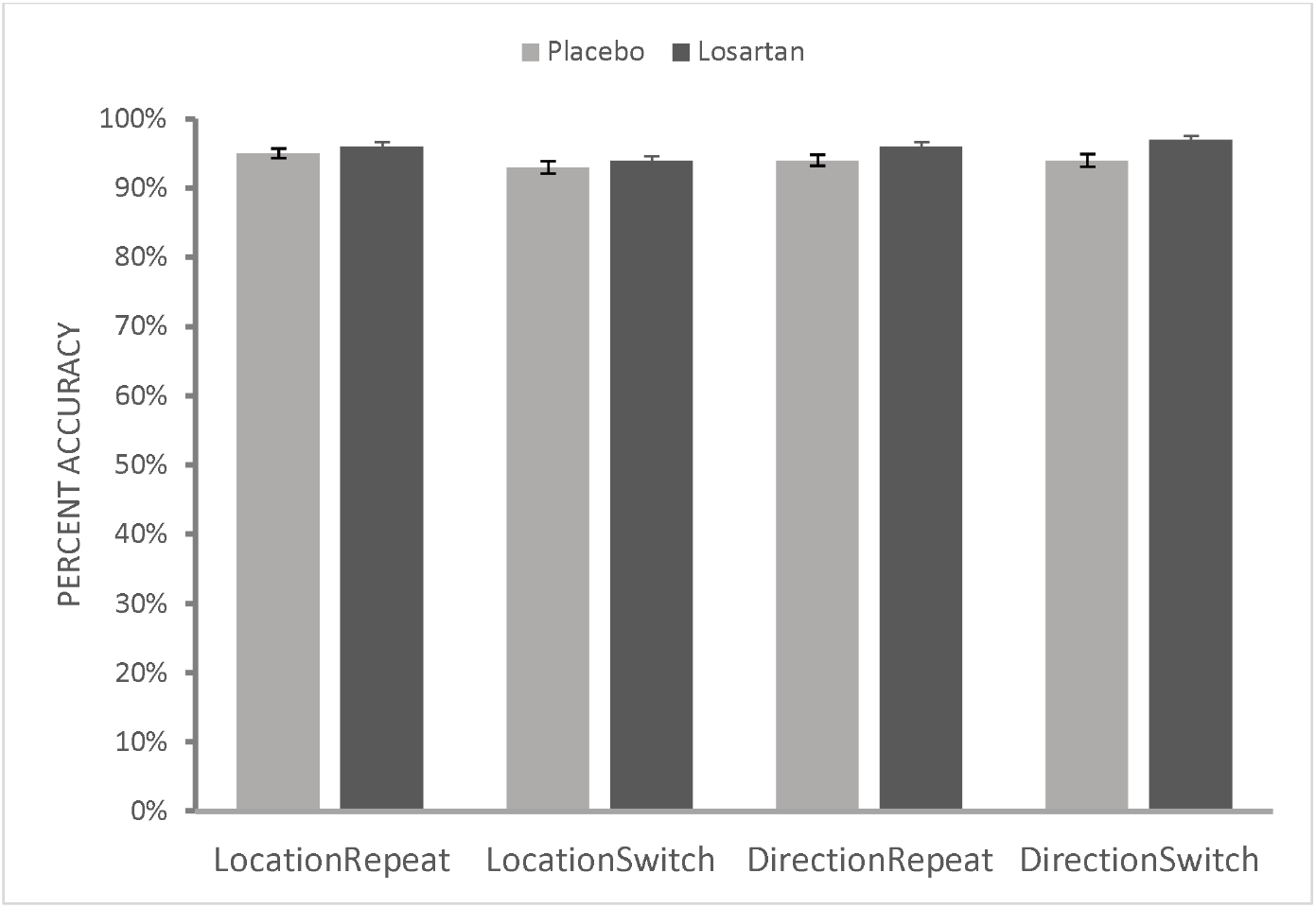
Mean percent accuracy across task and switch conditions between losartan and placebo groups.

### 3.4 Task-Switching as a Bin Score

When task-switching performance was assessed through a composite bin score, integrating RT and accuracy, there was no significant benefit of losartan (M= 1393.10, SD= 126.57) over placebo (M=1392.61, SD=150.13), (*t*(56) = 0.014, *p* = 0.99, Cohen’s d=0.04, see Figure 3).

**Figure 3:**
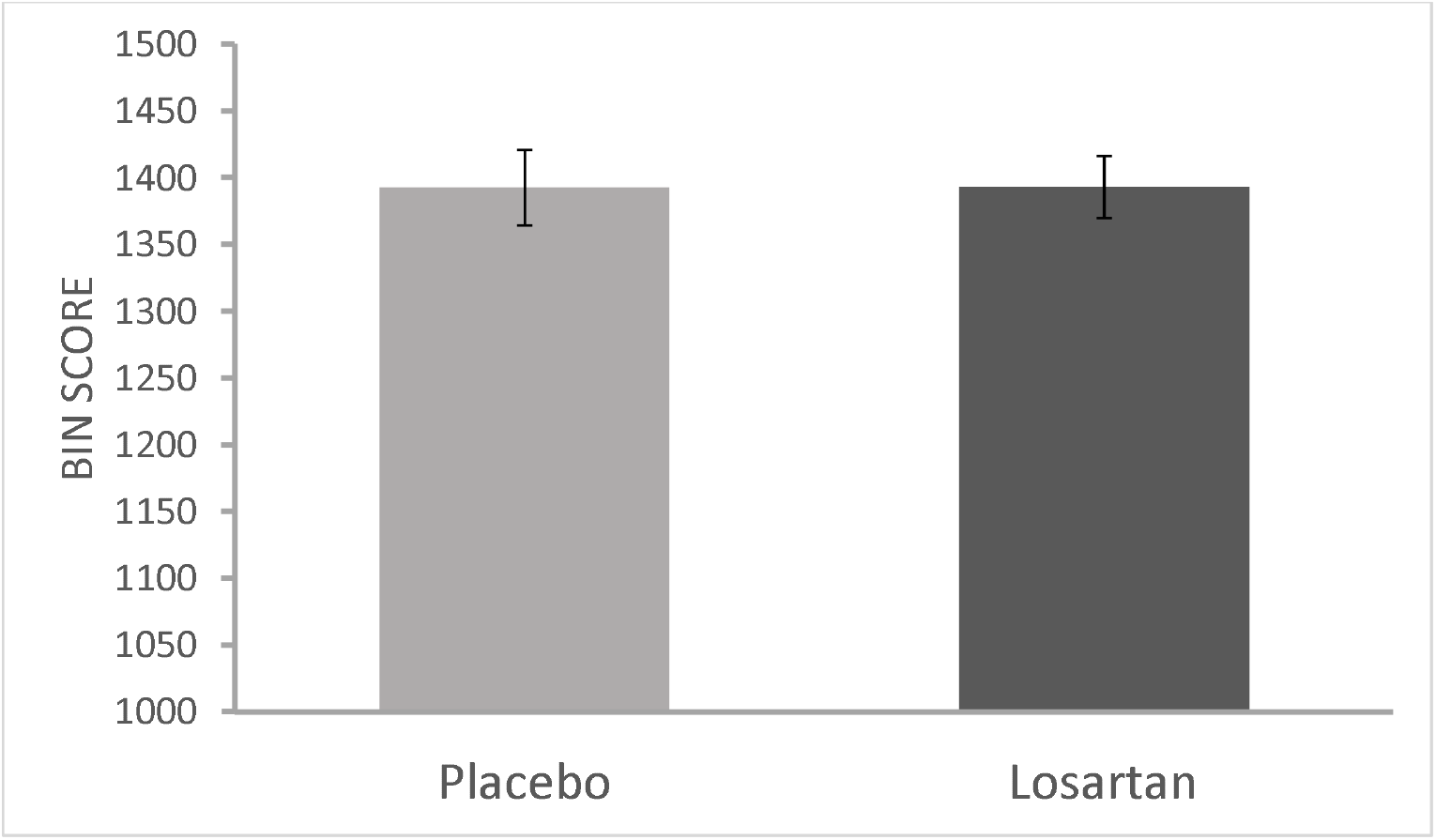
Mean composite bin scores, integrating RT and accuracy, for losartan and placebo groups.

## 4 Discussion

We assessed whether losartan, an angiotensin II receptor blocker, affects task-switching in healthy adults. Losartan did not significantly improve task-switching, although we observed consistently reduced RTs and higher accuracy levels in this group compared to placebo. When we analyzed group performance using a composite bin score, we again found no significant differences in task-switching between groups.

These findings are in contrast with our initial hypothesis—that losartan would benefit task-switching—and may be explained by several factors. First, it is possible that losartan’s effects were confounded by our healthy, high-performing population, where ceiling effects prevailed. Previous reports of losartan’s pro-cognitive effects (Mechaeil et al. 2011; Pulcu et al. 2019; Zhou et al. 2023; Prasad et al. 2025), coupled with the smaller switch costs observed in our study (placebo group: 11-37ms) compared to those seen in other young, non-clinical samples (33-56ms, Todorovic et al. 2022; 18-56ms, Gustavson et al. 2017) support this explanation. It is possible that because our participants were adept at task-switching, drug effects were less discernible than in populations with variable performance or cognitive impairment (e.g., Hajjar et al. 2022). Relevant to this, we found preliminary evidence that losartan improves direction switch trial accuracy, where a shift is required toward the harder, less prepotent direction task. This suggests that losartan may selectively impact cognitive flexibility when inhibition of a prepotent task is required and cognitive load is increased (Jost et al. 2017). This account aligns with losartan’s reported effects on other mechanisms that overlap with flexible thinking, such as fear extinction (Zhou et al. 2019), and its influence on regions implicated in cognitive control, such as the prefrontal cortex and striatum (Zhou et al. 2023; Bunge & Zelazo, 2006; Klanker et al. 2013). However, this exploratory finding is tentative, and it remains to be clarified whether RAS modulation on cognitive flexibility may be condition specific.

An alternative interpretation of our results is that losartan does not influence cognitive flexibility per se, but rather exerts modest effects on other aspects of cognition that affect task performance, such as processing speed, attention (Saxby et al. 2008), or motivation (Zhou et al. 2023). This explanation is compelling as it accounts for both the lack of switching effects and the numerically enhanced RTs and accuracy in the losartan group. Future studies may explore whether the RAS directly influences cognitive flexibility, cognitive processes that contribute to flexibility, or both.

The present study is characterized by several strengths. It is a novel, systematic investigation of RAS influence on cognitive flexibility in healthy individuals. Further, we applied a task-based measurement of a relatively sophisticated form of cognitive flexibility, task-switching. However, there are limitations. Our sample was composed of mainly young adults who were highly educated, impacting generalizability; further, we did not collect data surrounding race/ethnicity, which may have impacted drug effects (Helmer et al. 2018). We also only administered a single dose of losartan to healthy participants, precluding any conclusions about how this drug may affect task-switching over repeated administration or in clinical populations. Lastly, our task featured neutral stimuli, which may hold less ecological validity for psychiatric disease (Murphy et al. 2012).

## Conclusion

In summary, our findings suggest that losartan does not significantly impact task-switching in healthy adults, although these effects may have been difficult to detect in our sample. Future research may clarify RAS influence on cognitive flexibility by using other ARBs, more diverse samples, and/or alternative paradigms, which could help assess the utility of this system as a therapeutic target for mental health.

## CRediT Authorship Statement

**Divya Prasad:** investigation, project administration, formal analysis, data curation, and writing – original draft. **Georgia Feltham:** investigation, project administration, writing—review and editing. **Riccardo De Giorgi and Sara Costi:** resources, supervision, and writing – review and editing. **Elaine Fox:** resources, writing – review and editing. **Amy Gillespie:** formal analysis, data curation, supervision, and methodology and writing – review and editing. **Andrea Reinecke:** conceptualization, funding acquisition, methodology, resources, supervision, and writing – review and editing.

