## Supplementary Material for "The renin angiotensin system in cognitive flexibility: Effects of single-dose losartan on task-switching in healthy volunteers"

**Supplementary Online Content**

This supplementary material has been provided by the authors to give readers additional information about their work.

**METHODS**

***Sample Characterisation***

Our sample was characterized using several psychological questionnaires, including the State-Trait Anxiety Inventory-Trait Version (Spielberger et al. 1970), the Beck Depression Inventory (Beck et al. 1996), the Attentional Control Scale (Derryberry & Reed, 2002) the Intolerance of Uncertainty Scale (Freeston et al. 1994), the Perceived Stress Scale (Cohen et al. 1983), and the Temporal Experience of Pleasure Scale (Gard et al. 2006). Groups were comparable on all clinical characteristics (see Table 1 in main text).

***Randomization***

This experimental medicine trial utilized a 1:1 allocation ratio based on a randomization plan drawn up prior to study commencement by an external researcher uninvolved in participant testing. Specifically, study treatments were randomly allocated to sequential numbers for males and females (i.e., M1, M2, M3 and F1, F2, F3) in blocks of four. This meant that out of the first 4 female participants, two were given placebo and two were given losartan. There were thus 6 patterns for filling each block of 4 (i.e., using “L” for losartan and “P” for placebo: LLPP, LPPA, LPLP, PPLL, PLLP, PLPL). Each pattern was numbered 1 through 6 and a website ([www.random.org](http://www.random.org)) was used to create a random list of integers. Thus, the external researcher assigned this list sequentially to volunteers that entered the study.

***Data Cleaning***

Task-switching data was processed according to Wilcox and Keselman (2003), as followed by Gustavson et al. (2017) and Todorovic et al. (2022). Specifically: first, the initial four trials and trials with RTs below 150 ms were excluded. Data from two outlier participants (1 placebo, 1 losartan) were removed for low accuracy (<80%), indicating a possible failure to understand task instructions. For RT analyses, error trials and trials immediately after error trials were removed, in line with other task-switching protocols (Meiran, Chorev, & Sapir, 2000; Rogers & Monsell, 1995). Then, to calculate RT means for each participant, trials with RTs more than 3.32 times the mean absolute deviation were excluded from each participant’s data (Wilcox and Keselman, 2003; Gustavson et al. 2017). For participants with extremely high or low mean RTs, limits were set at 3 standard deviations from the group mean and outlier data were replaced with the nearest limit. All exclusions were determined with the assistance of a blinded researcher.

**RESULTS**

|  | **Location Repeat** | **Location Switch** | **Location Switch Cost** | **Direction Repeat** | **Direction Switch** | **Direction Switch Cost** |
| --- | --- | --- | --- | --- | --- | --- |
| ***RT (ms)*** |  |  |  |  |  |  |
| **Losartan** | 447.8 (91.5) | 476.3 (112.6) | 28.5 (50.1) | 551.1 (123.6) | 554.5 (132.4) | 3.3 (40.8) |
| **Placebo** | 480.2 (166.7) | 517.5 (168) | 37.4 (60.4) | 598.17 (168.5) | 610.1(201.6) | 11.5 (60.1) |
| **Total** | 463.4 (133.0) | 496.2 (142.3) | 32.8 (55.02) | 574.0(147.6) | 581.3 (170.1) | 7.3(50.8) |
| ***Accuracy (%)*** |  |  |  |  |  |  |
| **Losartan** | 0.96 (0.03) | 0.94 (0.03) | -0.018 (0.04) | 0.96 (0.03) | 0.97 (0.03) | 0.003 (0.03) |
| **Placebo** | 0.95 (0.04) | 0.93 (0.05) | -0.03 (0.03) | 0.94 (0.04) | 0.94 (0.05) | -0.001 (0.04) |
| **Total** | 0.96 (0.03) | 0.93 (0.04) | -0.02 (0.03) | 0.95 (0.04) | 0.96 (0.04) | 0.002 (0.03) |

***Table:*** *Mean RTs and accuracy for each trial type, alongside switch costs, by group and by total sample.*
